# Revealing the nervous system requirements of Alzheimer’s disease risk genes in *Drosophila*

**DOI:** 10.1101/2025.07.24.666383

**Authors:** Jennifer M Deger, Shabab B Hannan, Mingxue Gu, Colleen E Strohlein, Lindsey D Goodman, Sasidhar Pasupuleti, Zahid Shaik, Liwen Ma, Yarong Li, Jiayang Li, Morgan C Stephens, Michal Tyrlík, Zhandong Liu, Ismael Al-Ramahi, Juan Botas, Chad A Shaw, Oguz Kanca, Hugo J Bellen, Joshua M Shulman

## Abstract

Most Alzheimer’s disease (AD) susceptibility genes have poorly understood roles in the central nervous system (CNS). To address this gap, we systematically characterized 100 conserved candidate AD risk genes using a cross-species strategy in the fruit fly, *Drosophila melanogaster*. Genes were prioritized based primarily on human functional genomic evidence. We generated custom, loss-of-function alleles for each of the conserved fly orthologs. Most of the genes (80%) are expressed in the adult brain, including 24 neuron- and 13 glia-specific expression patterns. Overall, we identify 50 candidate AD risk gene homologs with requirements for CNS structure or function, including 18 whose loss of function causes neurodegeneration (e.g., *Snx6/SNX32* and *ClC-a/CLCN1*), 35 required for neurophysiology (e.g., *Arr1*/*ARRB2, stai/STMN4*), and 8 with diminished CNS resilience following a thermal or mechanical stress (e.g., *cindr/CD2AP*, *Amph/BIN1*). In a parallel screen, we found 28 AD risk gene homologs (e.g, *Ets98B/SPI1*, *Yod1/YOD1*) that modify the neurotoxicity of either amyloid-β peptide or tau protein, which aggregate to form AD pathology. To translate our findings back to human AD, we developed and deployed oligogenic risk scores based on gene clusters with shared nervous system phenotypes in flies, pinpointing functional pathways that differentially drive AD risk. Our results—available online via the Alzheimer’s Locus Integrative Cross-species Explorer (alice.nrihub.org)—reveal novel nervous system requirements for dozens of AD risk genes and may enable dissection of causal heterogeneity in AD.

## Introduction

Rapid advances in human genetics are revealing the major determinants of Alzheimer’s disease (AD) susceptibility. Initial studies of families with early-onset, autosomal dominant AD led to the discovery of *amyloid precursor protein* (*APP*)^1–3^ and *presenilin-1/2* (*PSEN1/2*)^4,5^, and genome sequencing studies in large population-based AD cohorts have begun to identify other rare, non-synonymous risk variants^6,7^. In addition, genome-wide association studies (GWAS) have identified many common variant associations, nominating hundreds of candidate genes as potential risk factors^8–11^. Whereas rare variants affecting protein function easily pinpoint the responsible genes, GWAS usually identify loci with many candidates. Interrogating GWAS loci using available large-scale human reference transcriptomes, proteomes, and epigenomes facilitates prioritization of likely causal genes^12–14^. For example, many AD susceptibility signals have been identified as expression quantitative trait loci (eQTLs), whereby genetic variants influence cell-type specific gene expression in microglia, astrocytes, and/or neurons^15^. Nevertheless, prioritizing AD risk genes based on human functional genomic evidence alone is limited by the available datasets. Further, many genes are broadly expressed in more than one cell type, requiring additional investigation to confirm the relevant cell type(s) and mechanism(s).

While progress in AD genetics has the potential to reveal novel therapeutic targets, most emerging candidate risk genes have unknown functions, especially within the nervous system. Moreover, the sheer number of emerging candidates creates a bottleneck to experimentally follow-up large numbers of genes, define their functions, and organize them into coherent pathways. Numerous AD experimental models are available, and many have proven powerful tools for mechanistic dissection of AD pathophysiology^16^. While it is possible to investigate many genes/variants in mammalian models, they are less feasible for systematic assessment of hundreds of candidates. Further, while cell-based screening strategies show promise^17,18^, many core features of AD, including the essential role of aging and the interplay among different cell types, are difficult to faithfully recapitulate *ex vivo*. As an alternative, non-mammalian systems including the fruit fly, *Drosophila melanogaster*, have a rich history of providing fundamental insights in neuroscience, including the identification and functional genetic dissection of human disease mechanisms^17,16,19^. The fly and human genomes share substantial homology^17,18,20,22,20–23^, including strong conservation of *APP*, *microtubule associated protein tau* (*MAPT*), and the majority of candidate AD genes identified from GWAS and sequencing analyses to date. *Drosophila* also offers high-throughput genetic analysis, enabling large-scale functional screening. Unbiased, forward genetic screens in fruit flies have successfully discovered many conserved molecular pathways important for human health including Notch in development^24^, Toll-like receptor signaling in innate immunity^25^, and the RAS-MAPK cascade in cancer^26,27^. Phenotypic screens in *Drosophila* have also defined genes required for neuronal development and maintenance^28–31^. For example, mutations disrupting retinal neurophysiology, assayed with electroretinograms, are enriched for fly homologs of genes implicated in AD and related neurodegenerative disorders^32^.

To systematically dissect the requirements of AD genes in the aging central nervous system (CNS), we screened 100 *Drosophila* orthologs of candidate AD risk genes for loss-of-function phenotypes. We selected complementary assays to profile gene requirements for brain structure, function, and resilience to stress. In parallel, we tested all genes for interactions with neurotoxicity caused by amyloid-β peptide (Aβ) or tau protein, which aggregate to form plaque and tangle pathology in AD. We also characterized the cell type-specific expression patterns of these 100 genes in the adult brain. Our study discovers novel CNS roles for dozens of AD risk genes and further highlights major functional pathways underlying AD risk.

## Methods

### Prioritization of AD risk genes

In the summer of 2022, we considered 493 genes spanning 90 AD susceptibility loci^33–36^. Each locus is defined as the region in linkage disequilibrium (LD) with the top AD risk variant (r^2^ ≥ 0.5), in addition to coding genes within 500 kb of either end of the LD block^33^. These genes were annotated according to functional genomic evidence from human studies. Genes with evidence that the AD risk variant alters the expression of the gene at the RNA^13^ or protein^37^ level were prioritized for further investigation. We supplemented the screen with two independent groups of genes from within AD risk loci: those that have published experimental evidence for involvement in AD, and those with AD-associated rare coding variants that are highly penetrant causes of familial AD. Detailed evidence and references for all genes’ prioritization criteria can be found in **Table S2**.

### Selection of fly orthologs

For each human gene prioritized for screening, a fly ortholog was selected based on the *Drosophila* RNAi Screening Center Integrative Ortholog Prediction Tool (DIOPT), version 8^38^. Some genes have two or more orthologs with the same DIOPT score. For these genes, we included two of the best orthologs: *CG8908* and *ldd* for *ABCA7*, *Kul* and *kuz* for *ADAM10*, *CtsB* and *CtsL1* for *CTSH, CG3860* and *Osbp* for *OSBP*, *SA1* and *SA2* for *STAG3*. We also screened two functional analogs of *APOE*, *GLaz* and *NLaz*, as this gene is not conserved in flies^39^. For other genes, one fly ortholog encompasses the functions of multiple human genes: *AdamTS-A* for *ADAMTS1* and *ADAMTS4, Psn* for *PSEN1* and *PSEN2*. We also included two fly homologs for *C4A* and *C4B*: *Tep3* and *Tep4,* as these fly genes have the same DIOPT scores and similar biological roles to both *C4A* and *C4B.* Fly genes that were screened, their human homologs, as well as current DIOPT scores can be found in **Table S4**.

### Generation of mutants

We systematically generated strong loss-of-function alleles, *T2A-GAL4* or *Kozak-GAL4*, for every gene included in this study when such alleles were not already available^40,41^. T2A was used for genes with a suitable intron, leading to truncation of the transcript as well as expression of GAL4 in the same spatiotemporal pattern as the mutated gene^40^. For genes without a suitable intron, the entire coding region was excised and replaced with a *Kozak-GAL4* cassette, which also leads to GAL4 expression in the same pattern as the gene where the cassette is inserted^41^. These reagents create strong loss-of-function mutations and have been extensively used to characterize gene functions^42^. *T2A-GAL4* or *Kozak-GAL4* lines were crossed to deficiency lines to determine whether homozygous mutants are viable. For genes on the X chromosome, we crossed female virgins carrying deficiencies to *T2A-GAL4* or *Kozak-GAL4* males, because males from X-chromosome deficiency lines typically carry only a balancer chromosome and thus cannot pass on the deficiency. For genes that are not on the X chromosome, we crossed *T2A-GAL4* or *Kozak-GAL4* virgins to males with a deficiency line missing the gene of interest. For non-essential genes, where adult progeny with *Gene^T2A-GAL4/def^* or *Gene^Kozak-GAL4/def^* are viable, we used these mutants for screening. For essential genes, where no adult mutants are viable, we crossed *T2A-GAL4* or *Kozak-GAL4* with 2 independent RNAi lines to achieve strong loss of function. When using *T2A-GAL4* for knockdown, we preferentially selected UAS-RNAi inserted downstream of the *T2A-GAL4* insertion. If two such lines were available from the Bloomington *Drosophila* Stock Center, we used them. Otherwise, we used RNAi lines from the Vienna *Drosophila* Resource Center. Catalog numbers for each line used, the genetic background for each mutant (*white*^+^ vs. *white^-^*), and information about viability can be found in **Table S4**.

### Screen for modifiers of tau and Aβ

For genetic screening of tau and Aβ-induced progressive locomotor dysfunction, we used 287 alleles targeting 100 *Drosophila* genes (**Tables S10 and S11**). These alleles were crossed with previously characterized *UAS*-*tau*^43,44^ or *UAS*-*Aβ42*^45^ combined with the *elav^c155^*-*GAL4* driver^46^ to achieve pan-neuronal expression. These lines also contain a heat-shock-induced lethality mutation on the Y chromosome (P{hs-hid}) to increase the efficiency of virgin collection^47^, and a balancer carrying *tub*-*GAL80^ts^* to silence the disease-associated protein, permitting maintenance of the stock. Virgin females were collected from this stock and crossed to males from the 287 alleles targeting 100 *Drosophila* genes. Crosses were set up on standard molasses-based *Drosophila* media at 25℃ under ambient room lighting. The female progeny from each cross were collected within a 12-hour period, then raised on standard molasses-based *Drosophila* media at 23℃ (*tau*-expressing flies) or 25℃ (*Aβ42-*expressing flies) under ambient room lighting. All modifier gene manipulations were tested in heterozygosity. We used manipulations predicted to result in both loss and gain of function. RNAi, MiMICs, deletions, EMS mutants, mutants made with attP swap, and P-element excisions are predicted to result in loss of function. UAS-cDNA are predicted to result in gain of function. P-elements and PBacs with GAL4-responsive UAS regulatory sequences (P{EP}, P{EPgy2}, PBac{WH})^48–50^ can have gain-of-function activity if the transposons are oriented in the direction of transcription (facing 3’ end). For controls, we included both *elav-GAL4*>*tau* or *elav-GAL4>Aβ42* crossed to either *w^11-18^* or “scramble” RNAi control (V2691), from the Vienna *Drosophila* Resource Center).

The negative geotaxis climbing assay was performed using a custom robotic system (SRI International, available in the Automated Behavioral Core at the Jan and Dan Duncan Neurological Research Institute) as previously described^51^. The robotic instrumentation elicits negative geotaxis (i.e., upward climbing) by tapping *Drosophila* housed in 96-vial arrays. After three taps, video cameras record and track the movement of animals at a rate of 30 frames per second for 7.5 s. For each genotype, we collect 4 replicates of 10 females to be tested in parallel (biological replicates). Each trial is repeated five times (technical replicates) per trial day. Replicates are randomly assigned to positions throughout the plate and blinded to users throughout the duration of experiments. Flies are transferred into new food every day.

The automated, high-throughput system is capable of assaying 16 arrays (1536 total vials) in ∼3.5 hr. To transform video recordings into quantifiable data, individual *Drosophila* are treated as an ellipse, and the software deconvolutes the movement of individuals by calculating the angle and distance that each ellipse moves between frames. The results of this analysis are used to compute the average speed of the 10 females in each vial/time point. Software required to run and configure the automation and image/track the videos include Adept desktop, Video Savant, MatLab with Image Processing Toolkit and Statistics Toolkit, RSLogix (Rockwell Automation), and Ultraware (Rockwell Automation). Additional custom-designed software includes Assay Control –SRI graphical user interface for controlling the assay machine; Analysis software bundles: FastPhenoTrack (Vision Processing Software), TrackingServer (Data Management Software), ScoringServer (Behavior Scoring Software), and Trackviewer (Visual Tracking Viewing Software).

We typically assess climbing from 5 to 18 days post-eclosion in our initial locomotor screen. On average, 6 timepoints were assessed for the tau model (as it shows an earlier locomotor defect) and 9 time points for Aβ. We measure locomotion in *Drosophila* as the average speed at which animals move in each vial as a function of age and genotype using a nonlinear random mixed effects regression model. ANOVA is used to assess statistical significance followed by Bonferroni-Holm correction to determine adjusted p-values. Specifically, we look at differences in regression between genotypes, genotypes with time (additive effect, represented by a shift in the curve), and the interaction of genotype and time (interactive effect, represented by a change in the slope of the curve). We estimate the expected statistical power to detect differences by each of our models using a threshold for statistical significance (P<0.05). We also calculate area between curves to measure the difference in speed over time between experimental animals and controls. In all cases we compare to control flies expressing *tau* or *Aβ42*. All plots are reviewed to confirm that statistically significant results represent meaningful differences on visual inspection. To consider a gene as an experimentally validated modifier gene, we require consistent results from at least 2 independent allele strains. Data for all strains tested can be found in **Table S10** (Aβ) and **S11** (tau).

### Expression patterns in adult brains

*T2A-GAL4* or *Kozak-GAL4* flies were crossed to flies carrying *UAS-mCherry NLS*. Progeny were aged for 14 days, at which time the whole brains were dissected in cold PBS and fixed overnight at 4°C in 4% paraformaldehyde. Brains were then permeabilized in 2% Phosphate-Buffered Saline with Tween 20 (PBST) for 1 hour at 4°C. Primary antibodies used were: rabbit anti-mCherry (GTX128508 from GeneTex) diluted 1:100, rat anti-elav (7E8A10 from Developmental Studies Hybridoma Bank) diluted 1:500 or mouse anti-repo (8D12 from Developmental Studies Hybridoma Bank) diluted 1:500. Primary antibodies were diluted in 2% PBST + 5% Normal Goat Serum (NGS). Brains were incubated in primary solution at 4°C for 24 hours, then washed twice quickly before five 10-minute washes in PBS. Secondary antibodies (goat anti-rabbit Cy5 and goat anti-rat or goat anti-mouse Cy3) were diluted 1:200 in 2% PBST + 5% NGS. Brains were incubated in secondary solution at 4°C overnight. The next day, brains were washed twice quickly before five 10-minute washes in PBS. Brains were then mounted on glass slides in VECTASHIELD® Antifade Mounting Medium (H-1000-10 from Vector Laboratories) without DAPI. Images were captured using a Zeiss LSM 780 confocal microscope, with a 20X air objective and the ZEN software. Fiji^52^ was used for image processing and analysis. We blinded ourselves to genotype and examined each section in a Z-stack of the brain and looked for colocalization between the green channel (elav or repo to mark neurons or glia, respectively) and the red channel (nuclear mCherry) in the same focal plane. Colocalization between these channels in the same plane was considered an indication of the gene being expressed in that cell type. Brains were annotated as being expressed specifically in neurons, specifically in glia, both neurons and glia, or neither. Results for brain expression patterns can be found in **Table 1** and **Table S1**.

**Table 1:**
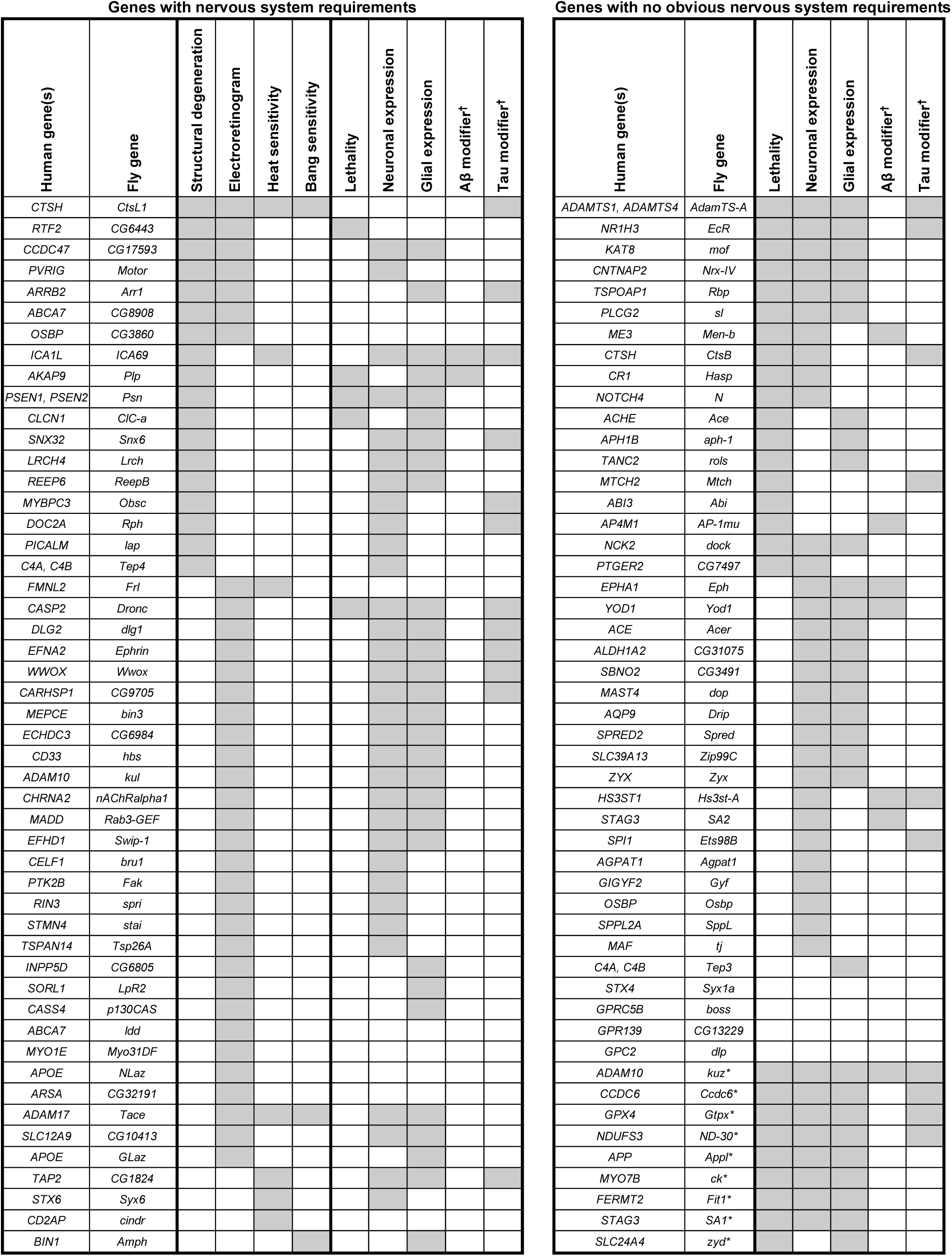
Summary of loss-of-function phenotypes for genes included in this screen. Each row represents a fly gene, and the corresponding human gene is listed in the first column. Genes whose disruption induces an effect as measured by brain histology (structure), electroretinograms (function), or heat/bang sensitivity (resilience) are shown on the left. Genes lacking nervous system phenotypes are shown on the right. *Loss of 9 genes was lethal even with partial knockdown and thus these mutants were not included for brain histology, electroretinograms, or stress sensitivity. **^†^**Only high-confidence tau/Aβ modifiers (with ≥2 independent strains showing the same direction of effect) are indicated in this table. Further information about all tau/Aβ modifiers can be found in Table S9.

### Whole-brain histology

Females were aged to 21 days at 25°C in ambient light. Proboscises, wings, and legs were removed before heads and thoraxes were placed in cold 8% glutaraldehyde and fixed for 7 days, rotating at 4°C. Samples were then transferred to plastic cassettes in 70% ethanol and brought to the Baylor College of Medicine Histology Core the same day for paraffin embedding. After embedding, two heads were sectioned and stained with hematoxylin and eosin. Sections were imaged and examined for vacuolar or other structural degenerative changes. We scored brain histology blinded to genotype. We counted the number of vacuoles in each brain. At least 2 heads were examined per genotype. We observed that wildtype control flies could have up to 10 vacuoles in the brain. Therefore, genotypes that consistently had ≥15 total vacuoles in the brain were annotated as having structural degeneration. For essential genes where two RNAi were used, both had to meet these criteria for the gene to be considered as required for maintenance of brain structure. Raw vacuole counts for all genotypes assayed can be found in **Table S5**.

### Electroretinograms

Flies were raised at 25°C in 24-hr light. Because absence of the *white* gene sensitizes photoreceptors to degeneration upon aging^53,54^, we tested white-eyed flies at 7 days, whereas we tested red-eyed flies at 21 days. Mutants that were created with *white* transgene-containing deficiency alleles or UAS-RNAi constructs had orange or red eyes and were compared to orange- or red-eyed controls. Mutants that were created without *white* transgene-containing alleles had white eyes and were compared to white-eyed controls. Details about genetic background and eye color for each mutant tested can be found in **Tables S1** and **S4**. Age-matched *y w* or *CantonS* flies were used as controls for white-eyed and red-eyed genotypes, respectively. To account for potential effects from genetic background, GAL4, and UAS, we also outcrossed *y w* virgins, which are in the same genetic background as *T2A-GAL4* and *Kozak-GAL4* mutants, to several strains, including a non-targeting RNAi control line from the Bloomington *Drosophila* Stock Center (B35785, *UAS-mCherry RNAi*) and an isogenic host strain from the Vienna *Drosophila* Resource Center RNAi library (V60200). All genotypes, including control genotypes, can be found in **Table S6**.

Electroretinogram recordings were performed blinded to genotype. Before recording, flies were gently fixed to a glass slide with Elmer’s glue. Recordings took place within one hour of gluing. A micropipette puller was used to pull fire-polished borosilicate glass capillaries (Sutter Instruments #BF120-69-10, outer diameter 1.2 mm, inner diameter 0.69 mm, length 10 cm) which were filled with 200 mM NaCl, inserted over silver wires, and used as recording electrodes. A grounding electrode was inserted into the head of the fly and a recording electrode was positioned on the surface of the eye. Recordings were performed in a Faraday cage to minimize background noise. Retinal voltage was recorded in response to 5+ light pulses with LabChart software (AD Instruments). 5 females were used per genotype, and 5 phototransduction traces were averaged per fly. The average of these 5 traces was used a single biological replicate. For each component of the phototransduction trace—light-coincident receptor potential (LCRP), on transients, off transients—statistical tests were performed with GraphPad Prism Version 10.5.0. Each genotype was compared with controls using a non-parametric one-way ANOVA, correcting for multiple comparisons with Dunn’s test. A significance value of P<0.05 was used as a cutoff to identify hits. For essential genes where two RNAi were used, both had to reach significance for the gene to be considered a hit. Data for all electroretinogram recordings can be found in **Table S6**.

### Heat and bang sensitivity

Flies were aged to 21 days at 25°C in ambient light. Males were typically used for this assay, but females were used for genes on the X chromosome (see “Generation of mutants”). We included *y w* or *CantonS* as wildtype controls. We observed that *y w* and *CantonS* strains were variably stress sensitive, so we also outcrossed *y w* virgins to males from several different genetic backgrounds, including *w^11,18^* males to match the genetic background of *T2A-GAL4* or *Kozak-GAL4* mutants that were crossed to *w^11,18^* deficiency lines. We also outcrossed *y w* virgins to *elav*-*GAL4* males to account for the potential toxicity of the GAL4, and to *CantonS* to account for potential effects of the *white* gene. Outcrossed lines were less stress sensitive than *y w* or *CantonS.* To ensure that the conditions to induce heat or bang sensitivity were met, we used *shibire^ts1^* used as positive control for heat shock, and *inaD^Δ174^* as positive control for bang sensitivity. Heat and bang sensitivity assays were performed blinded to genotype. For heat sensitivity, ∼10 flies were placed into empty plastic vials and submerged for 30 seconds in a 42°C water bath. For bang sensitivity, a vial containing ∼10 flies were vortexed at max speed for 30 seconds. Flies were given 10 seconds to recovery from each stress. Next, we recorded the number of flies impaired after the recovery period and calculated a percentage. Flies were considered impaired if they were supine and/or exhibiting seizure-like behaviors after the 10-second recovery period. For both heat and bang sensitivity, genotypes that were at least 60% impaired after exposure to the stressor were considered for follow up based on the primary screen. For essential genes where RNAi was used, both RNAi lines had to show at least 60% impairment. We selected 60% as the cutoff because this was more than double the average percentage impaired for controls across the screen (averages for all controls were 26% for heat sensitivity and 19% for bang sensitivity). For genotypes that were at least 60% impaired, we performed independent crosses and attempted to replicate the tests for heat and bang sensitivity, using 5-10 animals per biological replicate. Statistical tests were performed with GraphPad Prism Version 10.5.0. Each genotype was compared with controls using a non-parametric one-way ANOVA, correcting for multiple comparisons with Dunn’s test. A significance value of P<0.05 was used as a cutoff. Data can be found in **Table S7** (heat sensitivity) and **S8** (bang sensitivity).

### Human subjects and oligogenic risk scores

We leveraged clinical and genetic data from our previously described AD case cohort^55^. All studies of human subjects were conducted in accordance with the principles expressed in the Declaration of Helsinki. All patients with AD were diagnosed by neurologists with subspecialty training in memory disorders. The institutional review boards at Baylor College of Medicine and the Houston Methodist approved both the biospecimen collection and sequencing (H-50254; PRO00034371). The phenotypes included in this analysis were: decreased amplitude of depolarization from electroretinograms (CNS functional defects), brain vacuolar degeneration (CNS structural defects), bang/vortex sensitivity (CNS resilience), and heat sensitivity (CNS resilience). Each phenotype was coded as a binary categorial variable (present or absent). Each gene was mapped back to the relevant locus from GWAS and we used the AD risk allele for the lead single nucleotide polymorphism (SNP) based on the published source literature^11^. These data can be found in **Table S13**. For each phenotypic group, the number of risk alleles was tabulated for an independent cohort of n=99 AD cases using whole-genome sequencing data. The within-phenotype counts were then standardized by subtracting the mean risk allele count and dividing by the standard deviation. Next, a principal component analysis was performed on the standardized score.

The 2nd principal component was observed to differentiate individuals according to phenotypic class standardized scores, with loadings Z_ERG_= .35, Z_Structure_= .16 Z_Bang_= -.69 and Z_Heat_= -.61. This principal component explained 32% of the variation in the standardized risk allele count data. A sort order for individuals in the cohort was determined by computing the projection onto the 2nd principal component. For rendering, each individual’s scores were centered around their standardized score mean value. The heatmap was rendered using the R *ComplexHeatmap* package.

### Data presentation and software

Figures 4A, 6A, and 7A were created with BioRender.com. Figures 2 and 7B were created with R Studio version 4.5.0. Figures 5C, 6C, 6D, and S1 were created with GraphPad Prism version 10.5.0.

**Figure 1:**
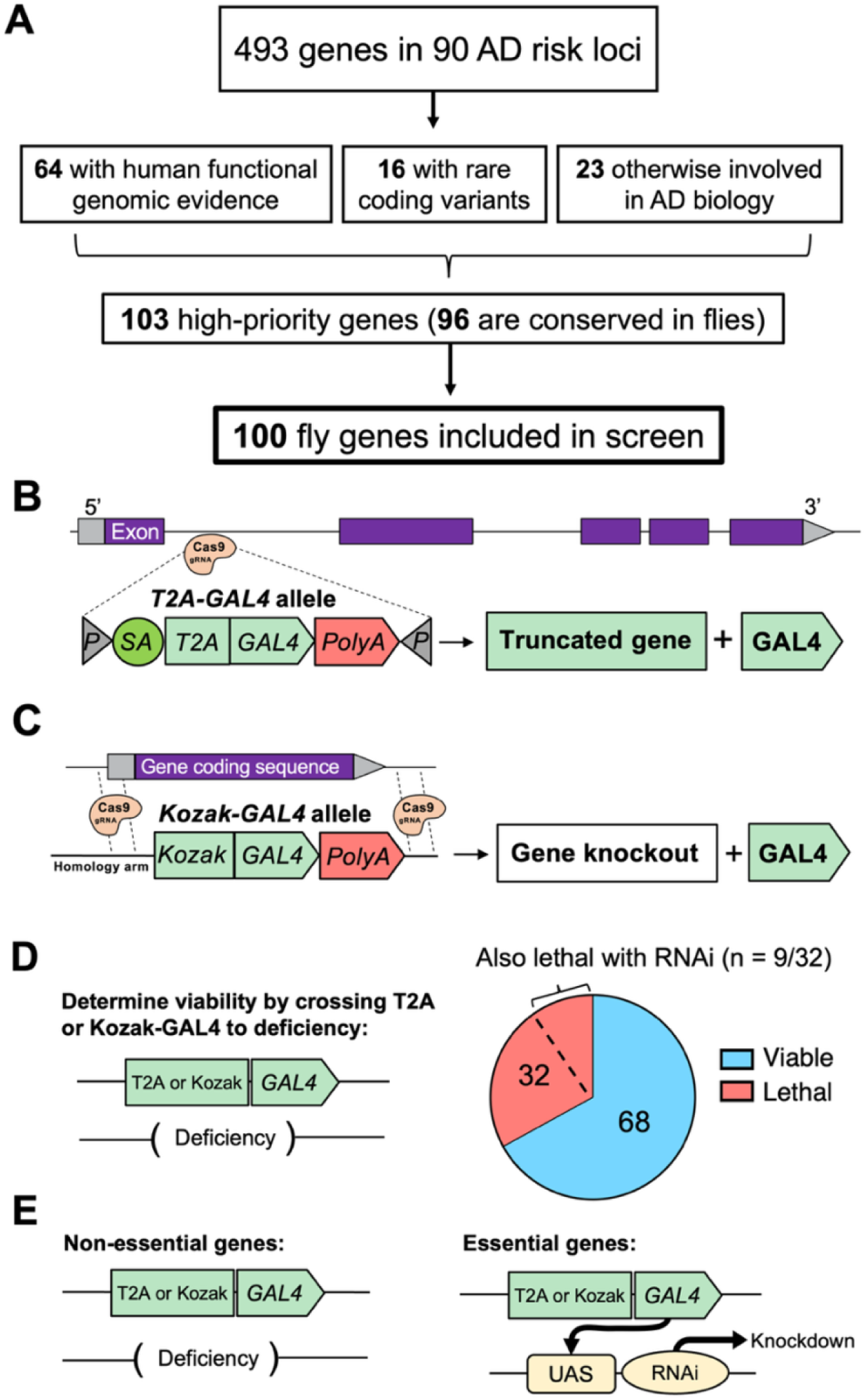
Strategy for AD risk gene prioritization and generation of custom loss-of-function alleles. A: 493 genes from within AD risk loci were initially considered and 103 were prioritized for screening. 96 of these high-priority genes are conserved in *Drosophila*. Because some human genes are orthologous to more than one fly gene, 100 fly genes total were included in the screen. B, C: Strategies used to create loss-of-function mutants (*T2A-GAL4* or *Kozak-GAL4*). D: *T2A-GAL4* or *Kozak-GAL4* alleles were crossed to deficiency lines missing the gene of interest to determine if double-copy loss of the gene was homozygous viable. 32 genes were found to be essential for adult viability. E: For non-essential genes, mutants for screening were created by crossing *T2A-GAL4* or *Kozak-GAL4* to deficiency lines. For essential genes, mutants were created by crossing *T2A-GAL4* or *Kozak-GAL4* to two independent UAS-RNAi lines. The RNAi strategy resulted in lethality for 9 genes. Detailed information about viability for each gene and strains used can be found in Table S4.

**Figure 2:**
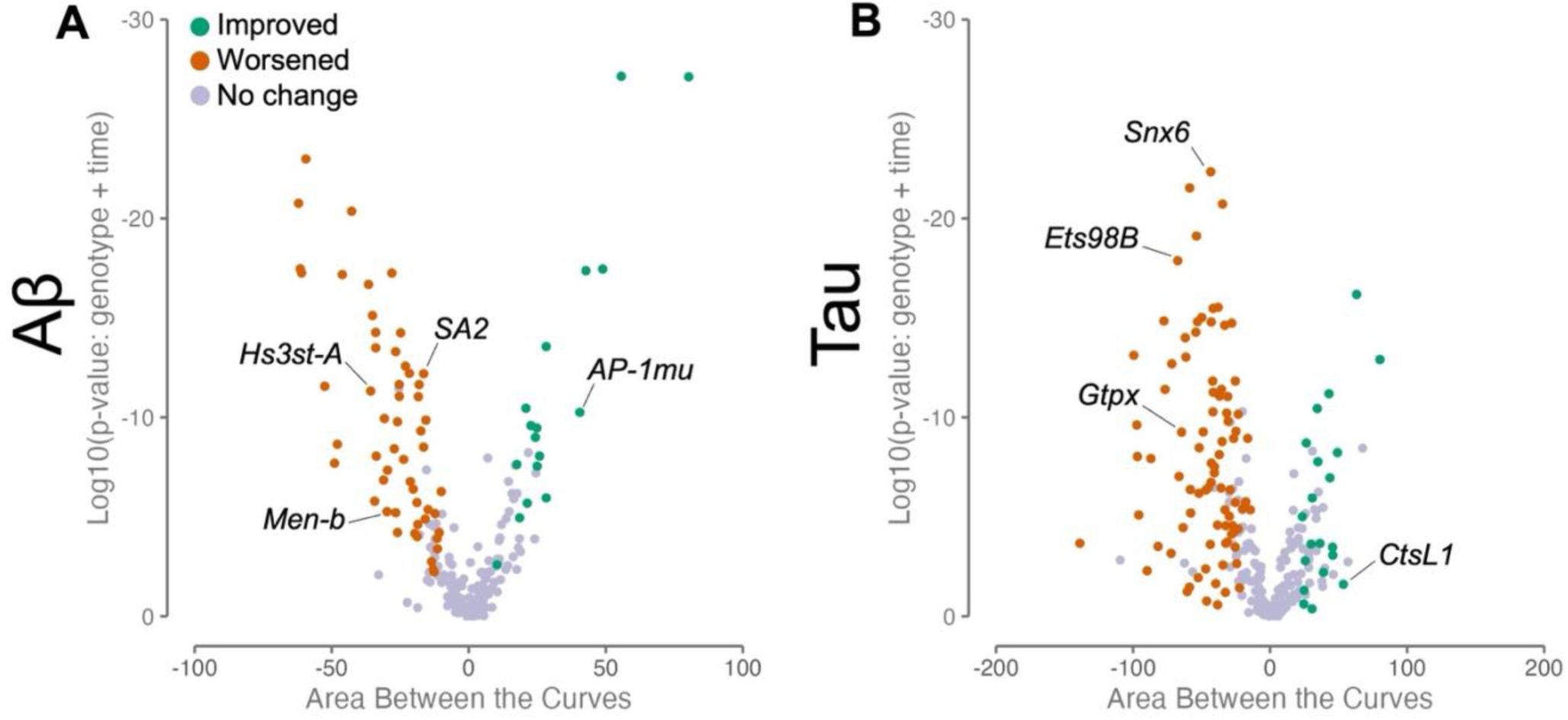
AD risk genes modify Aβ and tau-induced locomotor defects. A: Volcano plot for all strains tested in the Aβ model. The X axis represents the differences in areas between the speed over time for control vs. genetic manipulation. Log10(P) is shown on the Y axis. Each point represents a strain that was tested. If a strain worsens Aβ-induced locomotor defects according to a visual check of the climbing graph, it’s colored as orange. If a strain improves Aβ-induced locomotor defects, it’s colored as green. Strains with no effect are shown as purple. Selected examples are pointed out. Loss of *Hs3st-A/HS3ST1* (BDSC line 17884), *Men-b/ME3* (VDRC line 27535), and *SA2/STAG3* (V51569) worsens Aβ-induced climbing defects. Loss of *AP-1mu/AP4M1* (B27534) improves Aβ-induced climbing defects. B: Volcano plot for all strains tested in tau model, with modifiers denoted as in A. Loss of *Snx6*/*SNX32* (V110170), *Ets98B*/*SPI1* (B54520), and *Gtpx*/*GPX4* (B41879) worsens tau-induced climbing defects. Loss of *CtsL1/CTSH* (B5124) improves tau-induced climbing defects. A summary of results for all genes can be found in Table S9, and detailed information about each strain included can be found in Table S10 (Aβ) and S11 (tau).

## Results

### Prioritized candidate AD risk genes are conserved in *Drosophila*

To nominate AD candidate genes for further characterization, we first performed a systematic literature review, including results from the largest completed AD GWAS^8–11^ and complementary studies of rare genetic variants from either familial AD or case-control sequencing cohorts (Figure 1A and **Table S2**). Overall, we identified 493 genes spanning 90 AD susceptibility loci^33–36^ (**Table S3**). For common variant risk loci from GWAS, we included all candidate genes within a genomic window, considering local linkage disequilibrium in relation to the top AD-associated proxy variant (see Methods). For further prioritization, we next considered human functional genomic data, nominating 64 genes with previously published evidence that the AD risk variant is also associated with altered mRNA or protein expression in AD-relevant tissues, including data from brain, blood, other tissues^13,37^ (see **Table S2** for full list of references). Since there are gaps in available gene expression data and our knowledge of variant functional impacts is incomplete, we supplemented our list with 23 genes at susceptibility loci based on other published experimental evidence of links to AD mechanisms (**Table S2**). Lastly, we included 16 genes in which rare, non-synonymous variants are implicated, including established causes of autosomal dominant familial AD (e.g., *ABI3*, *ADAM10, APP*, *PSEN1/2*, and others^56–58^). In total, we prioritized 103 genes for loss-of-function screening, 96 of which are conserved in flies (Figure 1A). We identified 100 fly genes for further study (some human genes had 2 potential orthologs, e.g. *ADAM10* is orthologous to both *Kul* and *kuz* in flies). Based on FlyBase, a curated database of published data on *Drosophila* research^59^, 44 out of 100 (44%) conserved genes are not known to have a role in the CNS (**Table S12**).

### Conserved homologs of AD risk genes modify amyloid-beta or tau-induced CNS dysfunction

We next assessed whether the 100 fly orthologs of prioritized candidate AD risk genes may participate in Aβ- or tau-mediated disease mechanisms. Among our prioritized gene list, 18 out of 100 genes were previously screened for modifiers of either Aβ- or tau-induced neurotoxicity as part of a recently published study^60^, and the remaining genes were tested using the identical screening protocol as part of this work. Pan-neuronal expression of either the secreted human 42-amino-acid Aβ peptide or the tau protein (wild-type 2N4R isoform) recapitulates age-dependent neuronal loss and progressive CNS dysfunction, including locomotor impairment, and these experimental models have proven useful for prior functional dissection of AD candidate genes^60,61^. We used previously published fly strains^43,45^ harboring human transgenes responsive to the yeast Upstream Activating Sequence (UAS), whereby expression of *Aβ42* or *tau* is directed selectively to neurons by the *elav-*GAL4 pan-neuronal driver^62^. We obtained 287 independent genetic strains targeting the 100 *Drosophila* homologs, including RNA-interference (RNAi) transgenic lines, available loss-of-function alleles, as well as lines predicted to activate expression of the target gene **(Tables S10** and **S11**). We used an automated locomotor behavioral assay which is amenable for high-throughput genetic screening^60^. The climbing speed of adult flies was evaluated longitudinally and scored for worsening or improvement of the locomotor phenotypes caused by Aβ42- or tau-induced neuronal dysfunction (Figure 2). To minimize the possibility of off-target or genetic background effects, we considered genes as high-confidence modifiers when supported by consistent evidence from at least two independent strains or mutant alleles. Overall, 28 out of 100 genes evaluated were identified as high-confidence modifiers of either Aβ42-(n=9) or tau-induced neuronal dysfunction (n=22). 3 genes modified both Aβ42 and tau. Among the 28 modifier genes, the majority (n=24) are newly reported in this study (**Table S9**).

### Many homologs of AD risk genes are required for development and viability

Our initial studies establish that most AD candidate risk genes are conserved, and a substantial fraction may modify Aβ- and tau-mediated disease mechanisms. Nevertheless, most of these genes remain incompletely characterized with respect to their function in the nervous system. It is also possible that many genes function in pathways that are independent from Aβ and tau. To begin systematically characterizing gene functions, we generated loss-of-function mutations for each of the 100 fly gene orthologs of the prioritized candidate AD risk genes, using CRISPR genome editing to insert a *T2A-GAL4* (Figure 1B) or *Kozak-GAL4* (Figure 1C) cassette at each gene locus^40,41^. These lines are expected to truncate the transcript or delete the entire open reading frame, causing strong hypomorphic or null alleles, respectively. To determine whether the genes are essential during development, the *T2A-GAL4* or *Kozak-GAL4* alleles were crossed to available strains harboring chromosomal deficiencies predicted to delete the corresponding genes (Figure 1D). Overall, 32 out of 100 genes (32%) cause homozygous lethality, which is similar to genome-wide estimates for genes with essential developmental requirements in *Drosophila*^63^. For further studies interrogating the CNS requirements of non-essential genes (68 out of 100), we used the *T2A-GAL4* or *Kozak-GAL4* alleles in trans-heterozygosity with a deficiency allele (Figure 1E). This genotype is expected to behave as a strong loss-of-function allele combination (null or strong hypomorph) and minimizes potential confounding effects from second-site mutations, ensuring that the observed phenotypes are likely due to loss of the gene of interest.

For essential genes, we additionally crossed each *T2A-GAL4* or *Kozak-GAL4* to available RNAi transgenic lines to potentially create a viable, partial loss-of-function genotype for further investigation. In these flies, the GAL4 transcriptional activator is predicted to induce expression of the RNAi under the control of the endogenous gene regulatory sequences (Figure 1E). Even with this milder genetic perturbation, 9 genes remained lethal and were not pursued further for loss-of-function phenotyping (Figure 1D). However, we still characterized the expression pattern of these 9 genes in the brain and screened for modifiers of tau and Aβ-induced neurotoxicity. Information on viability and strain identifiers for each gene can be found in **Table S4**.

### Most homologs of AD risk genes are expressed in the adult brain

We next sought to systematically examine the brain expression of each prioritized, conserved AD candidate risk gene (Figure 3). We took advantage of the *T2A-GAL4* or *Kozak-GAL4* to drive expression of a genetically-encoded nuclear fluorescent reporter (*UAS-mCherry.NLS*) and co-stained for neuronal (anti-elav)^64^ or glial (anti-repo)^65^ nuclei. Most genes (98%) were expressed in the adult fly brain, including 24 with neuron-specific expression (e.g., *bru1/CELF1*, Figure 3A), 13 with glia-specific expression (e.g., *aph-1/APH1B*, Figure 3B), and 46 expressed in both brain cell types (e.g., *Psn/PSEN1* and *PSEN2*, Figure 3C). For 15 genes, we detected expression in adult brains but the signal did not clearly colocalize with neuronal or glial nuclear markers. For 2 genes (*CG32191*/*ARSA* and *CG3860*/*OSBP*), we did not detect brain expression using the mCherry reporter. These cases may be due to low expression levels or expression in other cell types. A summary of expression patterns can be found in **Tables 1** and **S1**, and representative images showing the adult brain expression patterns are also available in our custom online portal, the Alzheimer’s Locus Integrative Cross-species Explorer (alice.nrihub.org).

**Figure 3:**
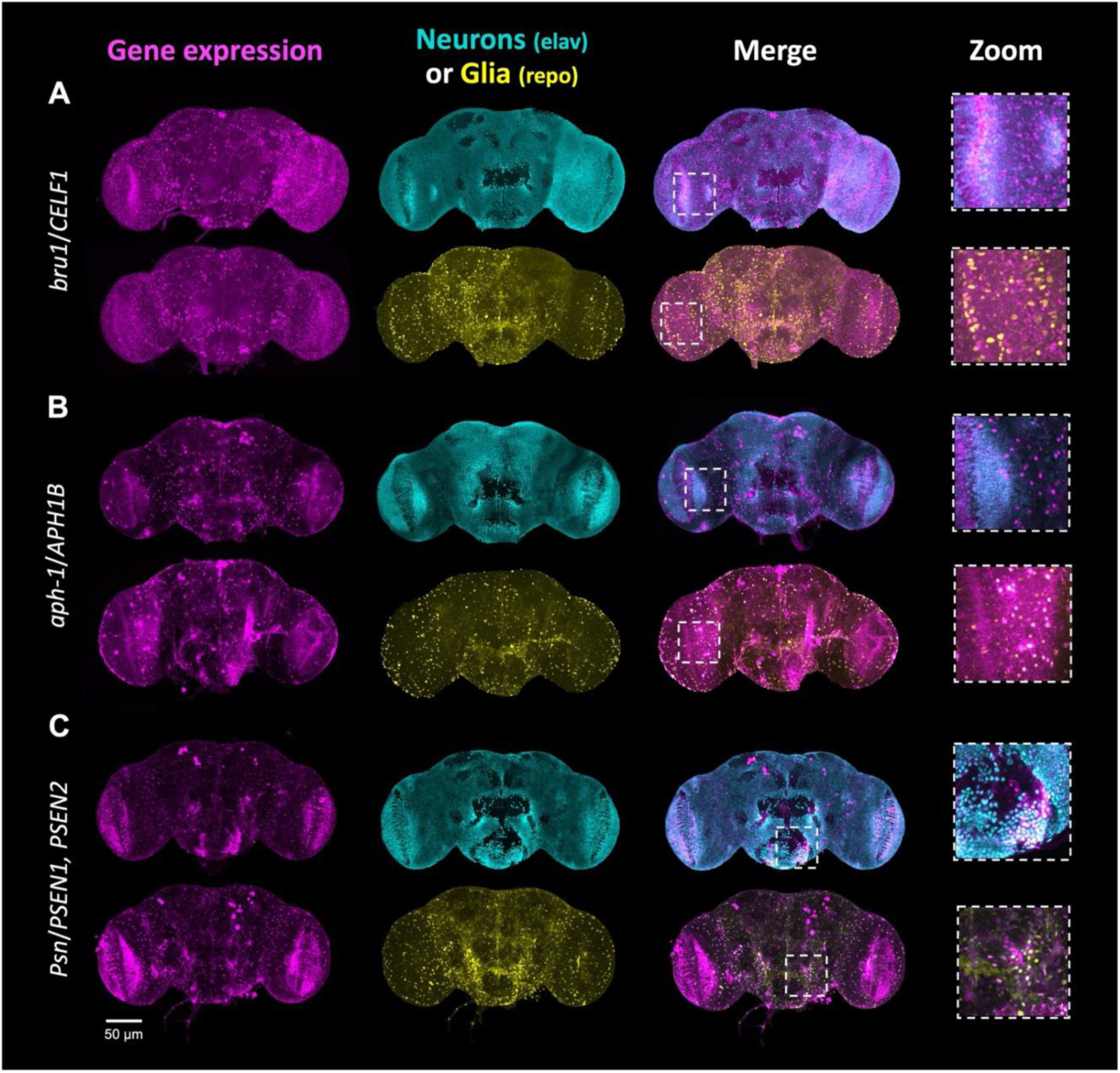
Cell type-specific gene expression patterns in adult brains. All 100 genes included in the screen were evaluated (see also Tables 1, S1, and the ALICE web portal: alice.nrihub.org). Examples of expression patterns (maximum intensity projections) are shown in A-C. Cells expressing gene of interest are shown in magenta (T2A-GAL4 or Kozak-GAL4 driving UAS*-*mCherry.NLS). An antibody specific to neurons (anti-elav) is shown in cyan, and an antibody specific to glia (anti-repo) is shown in yellow. Areas of colocalization appear as white. A: *bru1*, ortholog of human *CELF1*, has a predominantly neuronal expression pattern. B: *aph-1*, ortholog of human *APH1B*, is predominantly expressed in glia. C: *Psn*, ortholog of human *PSEN1* and *PSEN2*, is expressed in both neurons and glia. Scale bar = 50 μm.

### Several AD risk gene orthologs are required for maintenance of brain structure

To systematically examine the CNS requirements for each AD risk gene homolog, we next tested each of the CRISPR-based loss-of-function alleles (above) using several well-established assays for the study of neurodegeneration in *Drosophila*^66^. We first examined whether the homologs of AD candidate genes are required for the maintenance of brain structure (Figure 4). Mutations causing neurodegeneration in flies commonly cause the formation of large vacuoles within the neuropil^66^, and similar changes are also observed in transgenic fly models of AD in which human *Aβ42*^45,67^ or *tau*^68^ are expressed pan-neuronally. After aging flies to 21 days, we examined adult brain histology using hematoxylin and eosin-stained paraffin sections. Overall, we identified 18 genes causing significant structural degeneration following loss of function (Figure 4B**,C**). This group includes notable orthologs of AD gene candidates from GWAS (*lap/PICALM*), a proteome-wide association study (*Snx6/SNX32*), as well as a rare variant association analysis (*Plp/AKAP9*).

**Figure 4:**
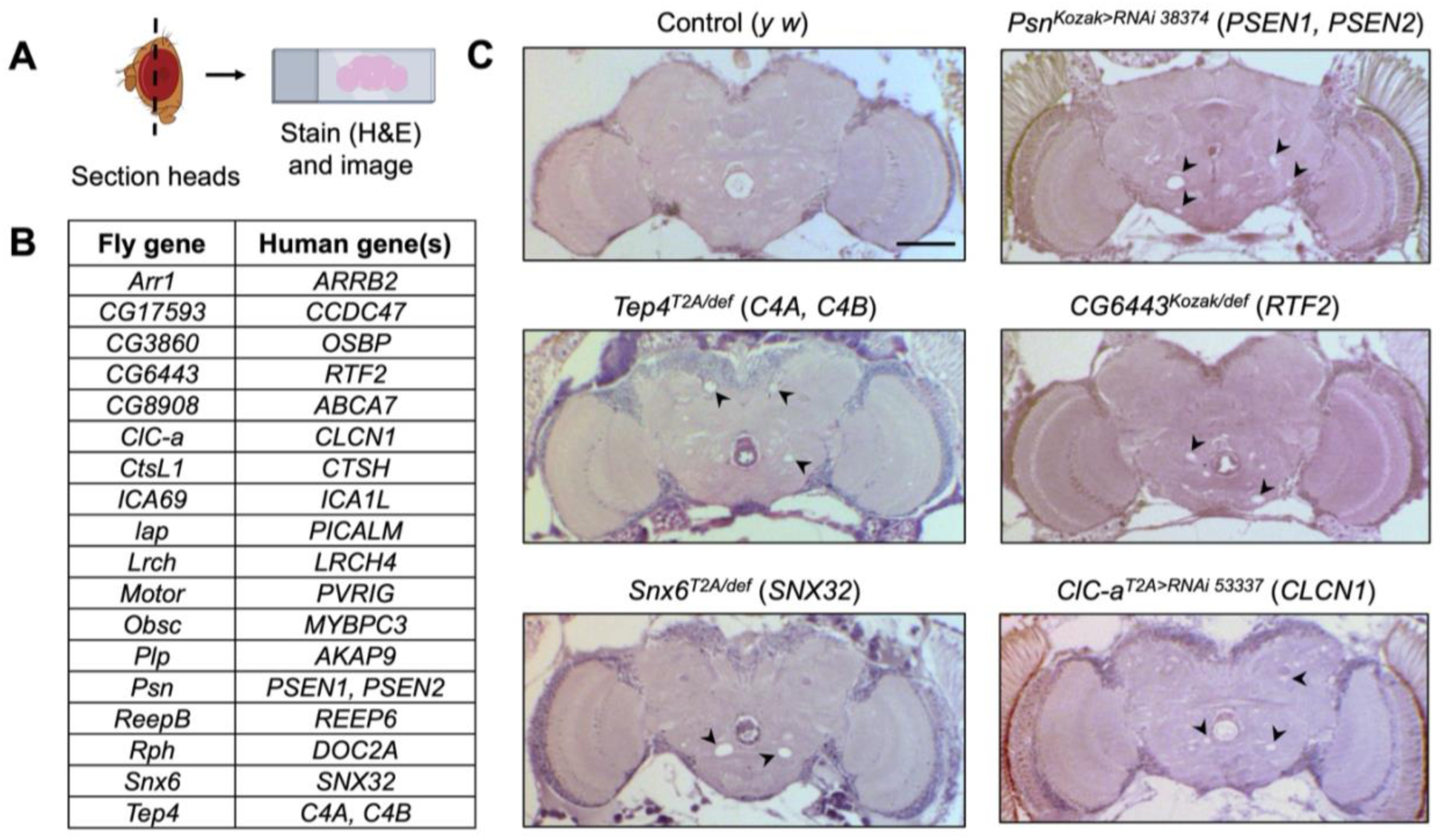
AD risk genes required for CNS structure based on adult brain histology. A: Experimental workflow. Female flies were raised to 21 days, then heads were dissected, fixed, and embedded in paraffin. Heads were sectioned, stained with hematoxylin and eosin, and imaged. B: List of the genes causing vacuolar neurodegenerative changes following loss of function (≥15 vacuoles in whole brain). D: Examples of genotypes with severe degeneration. Vacuoles are indicated with black arrowheads. The hole in the center of the brain is a normal feature. This is where the fly esophagus passes through. Scale bar = 50 μm. Vacuole counts for all genotypes can be found in Table S5.

### Loss of function in many AD risk gene orthologs disrupts neurophysiology

In AD, synapse loss and hippocampal circuit dysfunction precede neuronal loss and contribute to the earliest cognitive manifestations^69,70^. Therefore, we next turned to examining whether the orthologs of AD risk gene candidates are required for proper neurophysiology, including neuronal depolarization and synaptic transmission. We used the highly sensitive and quantitative electroretinogram assay, which is amenable to screening for *Drosophila* mutants with neurodegeneration^32^. Electroretinograms are extracellular recordings of retinal neuron response to light^71^. The resulting traces allow separate quantification of both photoreceptor depolarization (light coincident receptor potential or “amplitude”) as well as on and off transient potentials that represent postsynaptic responses to photoreceptor activity^72^ (Figure 5A).

**Figure 5:**
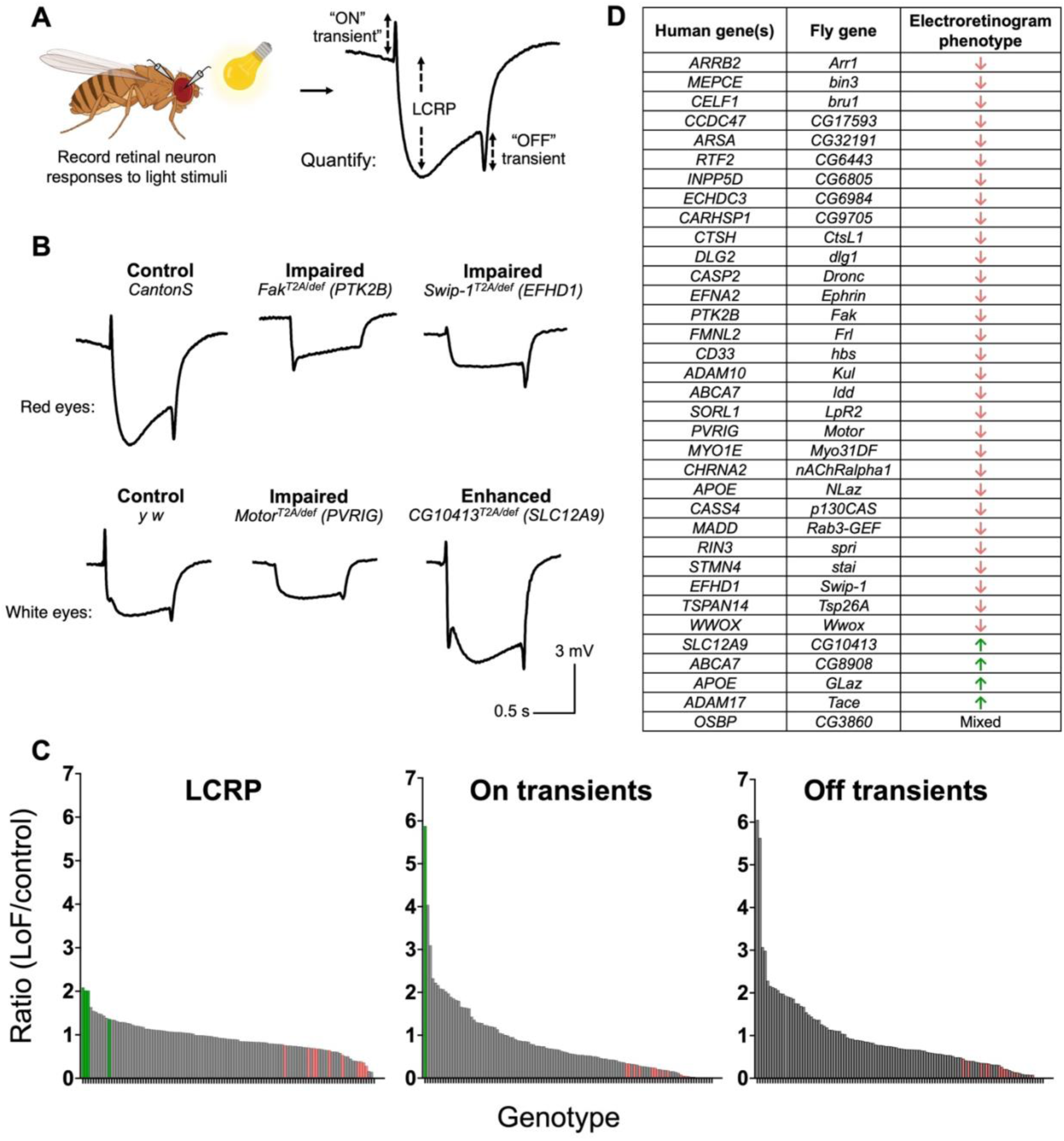
AD risk genes are required for neuronal function based on electroretinograms. A: Experimental workflow. Flies were aged in 24-hr light, then retinal responses to a light stimulus were recorded. An example of a normal trace and the three components of each trace are shown. LCRP= light coincident receptor potential. Created with BioRender.com. B: Examples of phototransduction traces of mutants with impaired or enhanced depolarization and transient potentials. Mutants were tested in both *white+* (red-eyes) and *white-* (white eyes) backgrounds. Thus, each mutant is compared to controls with matching eye colors. C: The averages for each component of the phototransduction trace are compared to controls from that day (ratio of mutant mV/control mV). Each bar represents one genotype. Those with statistically significant differences from controls are highlighted in green (increased mV) or red (decreased mV). D: Genes that had significant changes in one or more components of the phototransduction trace. Overall effect of gene loss-of-function is listed as decreased (red down arrow), increased (green up arrow), or mixed. Averages for all genotypes tested can be found in Table S6.

**Figure 6:**
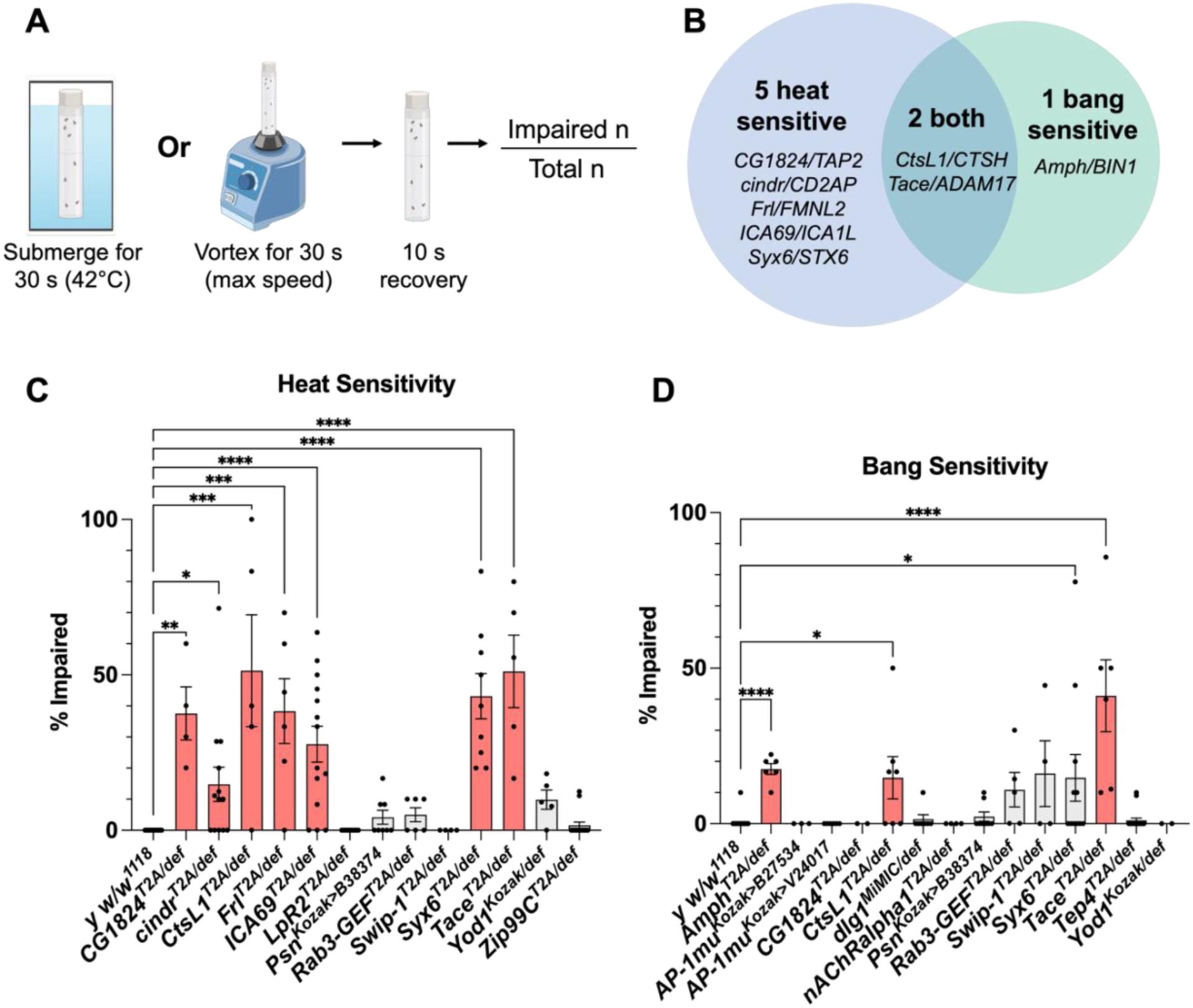
AD risk genes required for CNS resilience based on heat or bang sensitivity. A: Experimental workflow. Flies were aged to 21 days and then exposed to either a thermal stress (warm water) or mechanical stress (vortex) for 30 seconds. After a 10-second recovery period, the fraction of impaired flies was recorded. Created with BioRender.com. B: Venn diagram with the mutants that were heat sensitive, bang sensitive, or both. C: Quantification of replicate heat sensitivity assay, which was performed on genotypes that showed ≥60% heat sensitivity in the initial screen. D: Quantification of hits from replicate bang sensitivity assay, which was performed on genotypes that showed ≥60% bang sensitivity in the initial screen. Bars in C and D are SEM. Averages for all genotypes tested can be found in Tables S7-9.

**Figure 7:**
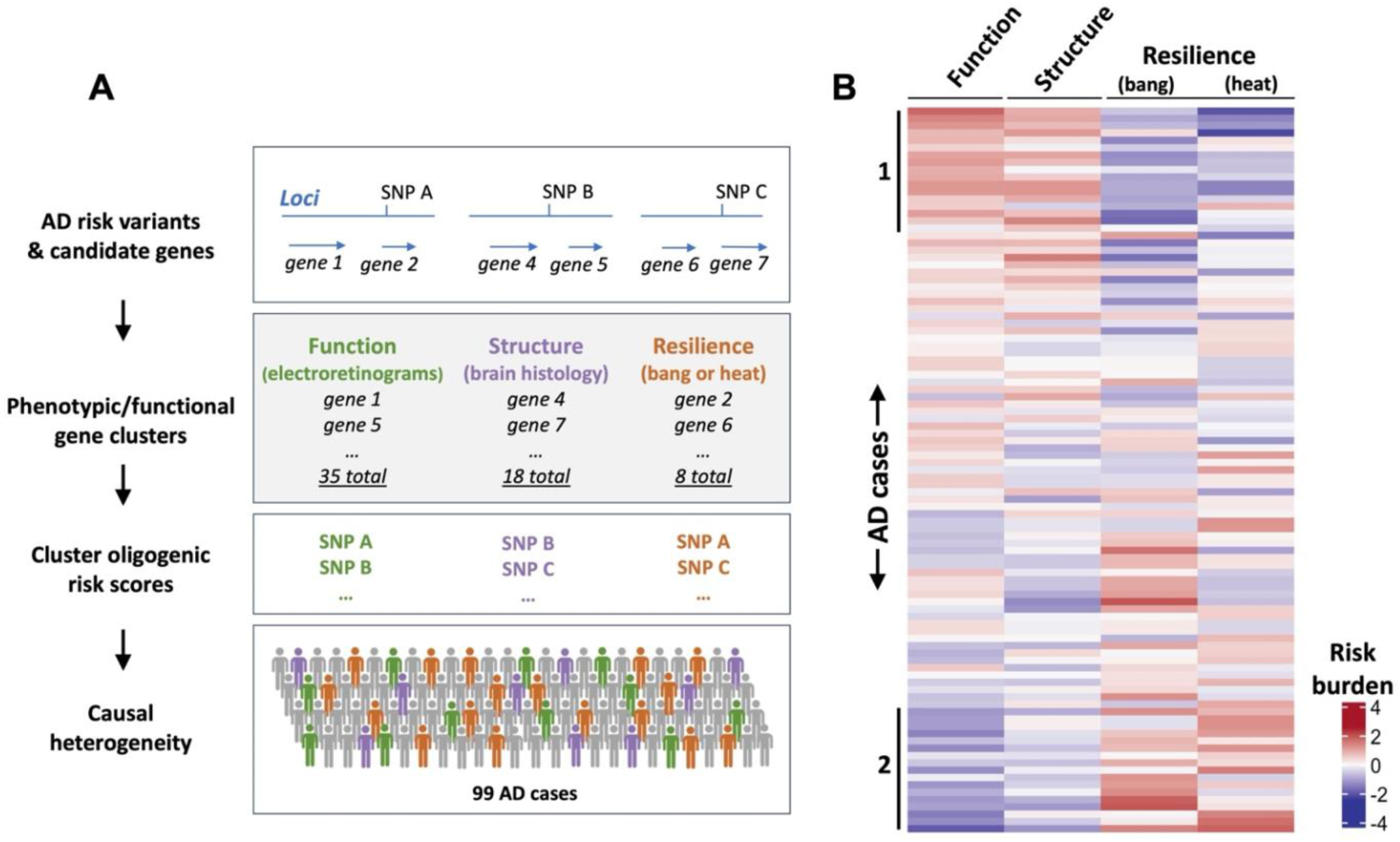
Cross-species dissection of AD causal heterogeneity. A: The lead AD risk variants were annotated for each locus based on published literature (see also Table S13). Phenotypic/functional gene clusters were then formed using the screening assays described in this manuscript. Genes are grouped based on shared requirements for neuronal function (electroretinograms), structure (brain histology), and/or resilience (bang or heat sensitivity). Multi-SNP oligogenic risk scores were derived from the marker variants linked to genes within each phenotype cluster. We computed scores highlighting the pathways driving AD risk among 99 individuals with AD. Created with BioRender.com. B: Heatmap showing oligogenic risk scores corresponding to the functional modules defined by the *Drosophila* screen. Each row represents a unique individual with AD. Red or blue color-scale denotes high-versus low-risk burden, respectively, for AD variants in genes with shared CNS requirements (columns).

We performed the neurophysiology assay in 2 genetic backgrounds: *white*^-^ (absent eye pigment)^73^ or wildtype (red eye color). The *white* gene, which controls eye pigment, has been shown to protect against retinal degeneration and can modify the electroretinogram response^53^. Hence, we consider *white^-^* a sensitized background for detecting genes with potential degenerative phenotypes^53,54^. We aged animals to either 7- or 21-days, for the *white^-^* or wildtype genetic backgrounds respectively, prior to performing the electroretinogram assay (see Methods). We counted determined which genotypes were significantly different from controls using a one-way ANOVA.

Overall, 35 AD risk gene homologs are associated with changes in retinal neurophysiology (**Table 1** and **Tables S6**). Loss-of-function alleles of 30 genes (e.g., *Swip-1/EFHD1, Fak/PTK2B, Arr1/ARRB2*) impair phototransduction, decreasing the depolarization amplitude, on transients, and/or off transients. By contrast, loss of function in 4 genes (*CG10413/SLC12A9, CG8908/ABCA7, Tace/ADAM17,* and *GLaz/APOE*) leads to a distinct phenotype, causing increased depolarization amplitude, indicating enhanced phototransduction. Interestingly, this second phenotype was only seen in flies lacking eye pigment, suggestive of a possible interaction with the sensitized *white^-^*genetic background. Lastly, mutation of 1 gene (*CG3860/OSBP*) produces a third, mixed phenotype in which retinal depolarization and on transients are increased but off transients are decreased. Example traces from mutants with impaired or enhanced phototransduction are shown in Figure 5B. An overview of all electroretinogram recordings is shown in Figure 5C. For genes to be classified as required for neuronal function based on this assay, we required robust and replicable results meeting statistical significance. The genes whose loss of function causes a significant effect in this assay are listed in Figure 5D.

### Many AD risk gene orthologs are required for neuronal resilience and recovery

Besides studying gene requirements to preserve brain structure and function, we also included assessments of CNS resilience and recovery. There is significant interindividual variation in the burden of AD pathology required for cognitive impairments, with many individuals showing relative resilience or vulnerability to similar degrees of pathology^74–76^. We therefore used 2 complementary assays, measuring the recovery of flies to either thermal stress (heat sensitivity) or mechanical stress (bang sensitivity). These assays are also sensitive tests of CNS hyperexcitability, which is increasingly recognized as important in AD pathogenesis^77–80^. The *Drosophila* heat and bang sensitivity assays have been previously used to study genes with roles in neurodegeneration^81,82^ as well other conditions with hyperexcitability, such as epilepsies^83,84^.

We subjected loss-of-function mutants for each conserved candidate AD risk gene to a 30-second submersion in a warm water bath or 30 seconds of vortexing and counted the percentage of flies that were impaired (supine and/or exhibiting seizure-like behaviors) after a 10-second recovery period (Figure 6A). We used a cutoff of 60% to identify potential hits, as 60% was well above the control average across the screen. We re-assayed mutants that were ≥60% sensitive with additional biological replicates to ensure robustness of our data, and determined which lines were significantly different from controls using a one-way ANOVA (Figure 6C, D). Overall, 8 genes were required for CNS resilience, causing impairments in recovery from either thermal or mechanical stress (Figure 6B). Specifically, 5 genes selectively increased heat sensitivity (e.g., *cindr/CD2AP*), 1 gene selectively increased bang sensitivity (*Amph/BIN1*), and 2 genes showed common requirements for recovery from both insults (*CtsL1/CTSH* and *Tace/ADAM17*). Data can be found in **Tables S7** (heat sensitivity) and **S8** (bang sensitivity).

### Genes with shared phenotypes define novel pathways underlying AD risk

Based on a century of *Drosophila* experimental genetics, it is well established that genes with shared loss-of-function phenotypes often function together^85,86^. This principle explains how fly genetic screens have successfully discovered the molecular mediators of many well-conserved signal transduction pathways (e.g., Notch, Wnt, RAS-MAPK) and mediators of the innate immune response^87^. We therefore reasoned that our systematic characterization of candidate AD risk gene homologs may pinpoint functional modules that mediate AD risk. Overall, we identify 50 genes with CNS requirements based on a positive result in at least one assay, including brain histology (18 genes), retinal neurophysiology (35 genes), and resilience (heat sensitivity: 7 genes; bang sensitivity: 3 genes) (**Table 1**). As expected, the phenotypes are partially overlapping, suggesting a genetic architecture comprised of both shared and unique features. For example, of the 30 genes with impaired retinal depolarization, 5 also caused brain histologic degeneration. Similarly, we noted a modest but significant negative correlation between retinal depolarization following gene loss of function and recovery time from either thermal or mechanical stress (**Figure S1**). Based on existing knowledge from the literature, gene clusters with shared phenotypes indeed appear to identify genes that may participate in cohesive functional modules. For example, among the 18 genes associated with vacuolar neurodegeneration, several are established regulators of endocytic trafficking (e.g. *lap/PICALM* and *Snx6/SNX32*)^88,89^ and others encode proteins that localize to the early endosome^90^. In addition, among the 4 genes causing increased phototransduction, 3 are known to participate in lipid metabolism (*CG8908/ABCA7, GLaz/APOE, Tace/ADAM17*)^90–92^.

AD is highly genetically heterogeneous, with at least 60 common variant susceptibility loci from GWAS as well as a growing number of rare variants risk alleles. We hypothesize that AD risk can be decomposed into discrete functional modules that may drive disease differentially among affected individuals. To test this hypothesis and translate our results, we leveraged the phenotypes identified from our cross-species analysis to partition AD causal heterogeneity. For genes that were prioritized from GWAS susceptibility loci, we mapped each gene back to the respective susceptibility locus and corresponding AD risk variant / SNP (Figure 7A, **Table S13**). In this manner, we derived multi-SNP oligogenic risk scores comprising the marker variants linked to genes within each phenotype cluster. The resulting 4 scores quantify the AD risk burden from common genetic variants linked to functional modules required for CNS structure, function, or resilience (heat or bang sensitivity). We computed scores for 99 locally recruited AD cases with genome sequencing and examined the relative risk burden corresponding to each of the gene-phenotype clusters (Figure 7B). Our results are consistent with causal heterogeneity among the functional pathways defined by our screen. For example, oligogenic risk scores based on genes required for retinal neurophysiology (function) and brain histology (structure) appeared positively correlated, identifying a subgroup of AD cases with elevated risk in both pathways (denoted as group 1 in Figure 7B). Interestingly, risk scores based on CNS resilience genes (causing heat or bang sensitivity) appear to be inversely correlated with the CNS structure/function scores, defining a distinct population subgroup in which AD risk may be driven by an alternate genetic pathway (group 2 in Figure 7B).

## Discussion

Despite remarkable progress in understanding the genetic architecture of AD, most candidate risk genes have poorly understood roles in the nervous system. This critical knowledge gap threatens to delay the translation of AD gene discoveries to new therapies. We have systematically examined the conserved homologs of 100 candidate AD risk genes for requirements in the fly CNS, generating valuable reagents, characterizing gene expression, and interrogating loss-of-function phenotypes. We discovered that loss of function in 50 genes—half of those prioritized for testing—disrupts brain structure, function, or resilience to stress. Based on FlyBase^59^, an online database of gene-phenotype relationships in *Drosophila*, 21 out of 50 genes were not previously known to cause nervous system phenotypes. For example, mutation of *Snx6*, encoding a potential regulator of protein trafficking, triggered vacuolar degeneration in the adult brain. The human homolog of *Snx6*, *SNX32*, was identified in an AD proteome-wide association study, which showed that the AD risk haplotype is associated with decreased *SNX32* protein expression^37^. Even among those genes that are already well-studied in the context of neurodegeneration, we discovered novel phenotypes suggestive of previously unknown functions. For example, whereas *lap* (homologous to human *PICALM*) has previously been linked to AD pathogenic mechanisms, such as synaptic vesicle recycling and lipid droplet formation^93,94^, we have newly discovered that *lap* loss of function also causes vacuolar degeneration on brain histology.

Rather than highlighting specific risk genes, GWAS usually identify genomic regions containing many candidates, creating a challenge to identify the responsible gene or genes from each implicated susceptibility locus^95,96^. Overall, our screen considered 493 genes from 90 discrete loci nominated from GWAS, with a mean of 5.4 genes per locus (range: 1-55 genes per locus, **Table S3**). Systematic experimental manipulations may be helpful for refining and fine mapping association signals to prioritize target genes. In instances where multiple genes were nominated and tested from a single locus, our results highlight CNS requirements for several genes in flies. For example, among 14 human candidate genes localizing to the *CLU/PTK2B* locus, 3 genes were prioritized for consideration in our loss-of-function screening, and orthologs of all of these, including *stai* (*STMN4* ortholog), *Fak* (*PTK2B*), and *nAChRalpha1* (*CHRNA2*), disrupted neurophysiology based on the electroretinogram assay. This result, along with similar findings from other susceptibility loci, may support the hypothesis that multiple genes at a given locus alter CNS function in combination. While gene selection and prioritization for our screen was strongly influenced by human functional genomic evidence, the available human expression datasets remain incomplete. Therefore, it is possible we have missed promising causal genes at susceptibility loci from among the 493 initially considered for testing. All the screening assays that we employed are high-throughput and readily scalable in the future for consideration of more comprehensive gene lists.

An important goal for our field is to understand how AD risk genes function coordinately within pathways. Literature-based curation of pathways has been one popular approach, highlighting enrichment of AD susceptibility gene candidates implicated in innate immunity, lipid metabolism, and endocytic trafficking^33,97^. However, knowledge gaps in the biomedical literature, including absent, incomplete, or even erroneous information on many genes, may limit the power and utility of this approach. Genetic screens in simple model organisms have a successful track record of identifying genes that work in the same biological pathways based on shared loss-of-function phenotypes^85,86,98–100^. Consistent with this, we found that genes whose homologs are associated with brain neurodegenerative phenotypes encode proteins that localize to the endosome (*ABCA7, PSEN1*) or regulate endocytic trafficking (*SNX32, PICALM, AP4M1*). Further, among 4 genes with homologs whose loss enhances neurotransmission, 3 (*ABCA7, APOE, ADAM17*) are implicated in lipid transport^90^ or lipid storage^101,102^.

Most individuals with AD coinherit multiple risk alleles that potentially act in combination. However, few studies have systematically attempted to dissect the underlying logic. If AD risk can indeed be broken down into discrete functional modules (i.e., structure, function, resilience), we might speculate broadly on two possible models. In the first, AD risk would be distributed evenly across the different pathways. In a second, contrasting model, AD susceptibility may instead be driven by a single, dominant pathway in each individual among the several alternative and heterogeneous functional modules. In our exploratory analysis, we partitioned AD risk due to common genetic variation into discrete functional clusters, based on loci with genes causing shared loss-of-function phenotypes in flies. Interestingly, we discover potential evidence supportive of both models. Specifically, we find that many individuals with AD in our cohort demonstrated a high cumulative burden of risk alleles from loci with genes required for brain structure (histology) and function (electroretinograms) or alternatively, from loci with genes required for resilience (heat/bang sensitivity).

In the future, it may be informative to expand the screening assays to include additional phenotypes. For example, many AD genes, including *ABCA7*, *APOE*, *PICALM*, and *CD2AP,* have been implicated in sequestering reactive oxygen species in glial lipid droplets, in part based on targeted screening in flies^39,94,103,104^. The *T2A-GAL4* / *Kozak-GAL4* reagents generated as part of this study will accelerate such future investigations. Since our primary objective was to understand the CNS requirements of AD genes, we relied on strong loss-of-function genotypes. In our complementary screen for modifiers of Aβ and tau-induced neurotoxicity, we employed a variety of additional alleles and genotypes that may better approximate the modest gene perturbations caused by AD-associated regulatory variants. Interestingly, among the 28 genes discovered to modify Aβ or tau, there was minimal enrichment for those genes with CNS phenotypes (**Table 1**). Reciprocally, the requirement for CNS structure, function, and/or resilience was not a strong predictor that a gene would also interact with Aβ or tau. Our results may be overall consistent with dose-sensitive, pleiotropic roles for AD genes in either supporting healthy brain aging or responding to neuropathology.

In sum, we have generated a wealth of experimental data and valuable reagents relevant to understanding the CNS functions of AD risk genes. All results are available via the Alzheimer’s Locus Integrative Cross-species Explorer (ALICE): alice.nrihub.org. The portal readily accepts searches of human or fly gene names for broad accessibility, and users are invited to nominate additional candidates for testing when inputting genes that have not yet been evaluated. We have employed scalable assays and can readily test additional genes to expand this resource in the future. Our findings reveal novel CNS requirements for many genes, define relevant genetic pathways, and begin to dissect causal heterogeneity for AD.

## Acknowledgements

This work was supported by National Institutes of Health grants, including T32GM136611 (JMD, CES, MT), F31NS129062 (MCS), R01AG074009 (IA), R01AG073260 (HJB), R24OD031447 (HJB, OK), and U01AG072439 (HJB, JMS, JB), T32NS043124 (LDG), P50HD103555, and U54HD083092. Additional support from foundations was provided by the Baylor Research Advocates for Student Scientists (JD, MT), The Robert and Janice McNair Foundation (CS), Southern Star Medical Foundation (JMS, OK), and the BrightFocus Foundation (Postdoctoral Fellowship Program in Alzheimer’s Disease Research to LDG, A2021008F). We’d also like to thank the Baylor College of Medicine Histology Core and the Intellectual and Developmental Disabilities Research Center. We would like to thank Drs. Shinya Yamamoto and Dr. David Li-Kroeger for helpful discussions of experimental design, and to the technicians of the Gene Disruption Project: Catherine Grace Burns, Megan Cooper, Ming Ge, Wen-Wen Lin, Junyan Fang, Ying Fang, Minhua Huang, Minghua Qin Idso, Hongling Pan, Jin Yue, Ruifang Zhang, and Xue Zheng for technical assistance.

## Author contributions

Conceptualization: JMD, HJB, JMS

Formal analysis: CAS, MT

Funding acquisition: HJB, JB, JMS

Investigation: JMD, SBH, MG, CES, LDG, LM, YL, JL, MS

Resources: OK

Software: SP, ZS

Supervision: IA-R, JB, ZL, HJB, JMS

Writing – original draft: JMD, HJB, JMS

Writing – review & editing: JMD, HJB, JMS, CAS, MT, SBH, MG, CES, LDG, LM, YL, JL, MS, OK, SP, ZS, IA-R, JB, ZL

## Declaration of interests

The authors have no relevant disclosures.

## Web resources

All results are available online via the Alzheimer’s Locus Integrative Cross-species Explorer (alice.nrihub.org).

---

**Figure S1:**
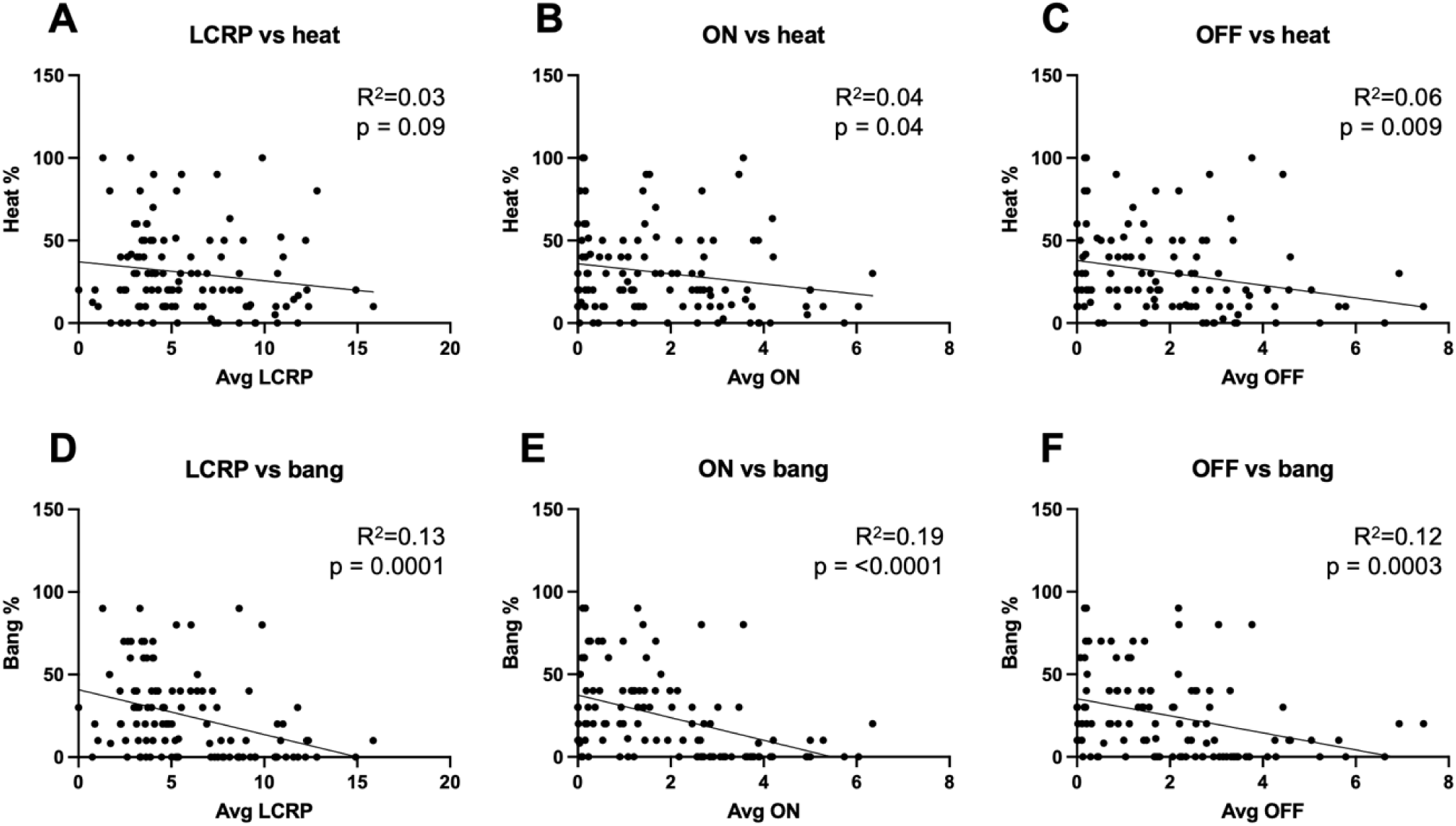
Negative correlation between electroretinograms and stress sensitivity. Each point represents a genotype. Average LCRP, on transient, or off transient values for each genotype are shown on the X axes. Percent impaired after heat or vortexing (bang) in the initial screen is shown on the Y axes. R^2^ and P values were calculated using simple linear regression.

## Notes

https://alice.nrihub.org/

### Competing Interest Statement

The authors have declared no competing interest.

